# Biofilm compactness and spatial community composition determine bacterial phage-susceptibility

**DOI:** 10.64898/2026.09.15.751744

**Authors:** Mads Frederik Hansen, Johannes Højlund Olsen, Agnete Nielsen Psilander, Amaru M. Djurhuus, Sigrid Lundgren Thomsen, Hannah Jeckel, Nanna Randløv Hartvig, Eva Jiménez-Siebert, Carey Nadell, Knut Drescher, Lars Hestbjerg Hansen, Mette Burmølle

## Abstract

Bacteriophages (phages) are viruses that infect bacteria, often posing a threat to their survival. In recent years, research has shifted from studying bacteria-phage interactions in homogeneous, well-mixed environments to investigating more complex heterogeneous, structured communities, thereby more accurately reflecting bacterial life in natural and disease-associated settings. So far, these studies predominantly tested interactions of phages and single-species biofilms, despite the dominance of highly diverse biofilm communities in nature. Here, we investigated how different lytic phages spread in and infect a synthetic community composed of *Escherichia coli*, *Kluyvera cryocrescens,* and *Vibrio anguillarum*. We found that *E. coli* was protected from eradication by the T7-coliphage when embedded in a multispecies biofilm and covered by the two other species, where the *V. anguillarum* biofilm matrix was essential for protection of *E. coli*. The protective nature of the *V. anguillarum* matrix also ensured survival of *K. cryocrescens* exposed to the lytic kluyveraphage Lyn. *V. anguillarum* mutants lacking the matrix synthesis genes *vpsMNOP* or *rbmC* failed to protect *E. coli* despite forming biofilms. Labelling of phages indicated that these were not trapped in the matrix of the wildtype. Instead, advanced image analysis revealed that the mutants displayed biofilms with lower cell density, resulting in the *E. coli* clusters being less tightly surrounded by *V. anguillarum* cells, which opened gaps for phage entry. These findings highlight that community context and matrix composition play a major role in phage-bacteria dynamics.

## Introduction

Many bacteria form dense matrix-embedded communities known as biofilms [1]. The matrix ensures spatial rigidity and, as the community expands, its physiochemical and metabolic landscape becomes spatially heterogenous [2]. The spatial variation of environmental conditions inside biofilms facilitates the coexistence of multiple species for extended periods of time [3,4]. In the community, the eco-evolutionary dynamics and physiological structure are governed by interspecies interactions [5–7]. In fact, there are multiple examples of community-intrinsic properties, where the community phenotype and architecture cannot be predicted by studying single species in isolation [8].

Different organisms exhibit different susceptibilities to a range of toxins and predators [9,10]. Bacteriophages (phages), which are viruses that infect and kill bacterial cells, typically have a narrow host range and only infect a fraction of strains of a given bacterial species [11]. Nonetheless, the abundance and diversity of phages provide a strong selective pressure in nature, as reflected in the wide arsenal of molecular phage defence systems discovered in recent years [12–15]. Protection against phages can also be facilitated by extracellular components [16]. Exopolysaccharides protect aggregates of *Pseudomonas aeruginosa* [17] and amyloid curli fibres in mature biofilms can shield *Escherichia coli* from infection as well [18]. Recently it was also shown that dual-species biofilms were able to provide community protection [19]. Specifically, *E. coli* embedded in a *Vibrio cholerae* biofilm was protected from killing by a coliphage. The protection was dependent on the degree of intermixing of the species, and the *E. coli* protection was significantly reduced in biofilms with a *V. cholerae* Δ*rbmA* mutant with less dense community architecture [19]. These findings highlight the protective nature of biofilms but also indicate that certain features of the biofilm architecture, such as cell density, are pivotal for the kinetics of phage infections. Since polymicrobial communities impact agricultural and industrial production [20] and are of clinical relevance [21,22], a deeper understanding of how such communities withstand phage exposure and how phages change community dynamics may be instrumental for ecological and industrial applications, e.g. for targeted microbiota engineering [23,24].

Here, we studied the impact of matrix composition and architecture on bacteria-phage interactions in a three-species biofilm community. Specifically, we used a biofilm community composed of *E. coli*, *Kluyvera cryocrescens* and *Vibrio anguillarum* with each species manipulated to encode a respective fluorescent reporter, enabling spatial and temporal quantification of population dynamics. In contrast to *V. cholerae*, for which biofilm regulation and components are well-studied [25], the specific formation mechanisms and components of *V. anguillarum* biofilms are unexplored. When exposing this community to different phages, we observed phage protection dependent on the presence of *V. anguillarum* and revealed that it was conditional on the presence of specific biofilm matrix components and their consequence on the biofilm architecture. Notably, the biofilm phenotypes of *V. anguillarum* matrix mutants were different from what has previously been reported for *V. cholerae*. For example, the strain used here was able to protect *E. coli* from phage killing despite not encoding *rbmA*, which was found essential for dense *V. cholerae* biofilm-facilitated protection [19]. In summary, our study shows that protection from phage infection in multispecies communities is determined by biofilm architecture and spatial organization and that different matrix components contribute to this protection in different species.

## Methods

### Biofilm glass transfer assay

Biofilm communities were established on round cover glasses in the wells of a 12-well plate (Fig. 1). To elevate glass slides from the bottom, they were placed on a single layer bed of Ø4mm glass beads. For inoculation, overnight cultures of bacteria were diluted 100x in Lysogeny Broth (LB) Miller and incubated with shake at 30°C for 2h, then diluted to OD_600 =_ 0.01 from where 1.5ml were added to the wells of choice. Plates were then incubated statically at 30°C. After 24h, a sterile tweezer was used to move the glass slides to new wells with fresh LB Miller or a bacterial culture of a different species prepared as described for initial inoculation. After 48h of incubation, the glass slides were transferred once more to either fresh media or to a well with the third bacterial species. After 72h, slides were moved to a well with fresh medium in which phages (coliphage T7 or Kluyveraphage Lyn) were added to a final concentration of 2.25×10^7^ PFU/ml from a high titer stock (approx. 2×10^10^ PFU/ml). Glass slides were never inverted during transfer between wells. Biofilms were cultivated with phages for 18h at 30°C before glass slides were imaged by inverting the slide and placing it on a larger rectangular cover glass, with an agar pad on top to decrease desiccation while imaging on an inverted microscope. Consequently, the side of the glass slide facing upwards during incubation was the one imaged. Reagents, bacterial strains and phages are listed in Suppl. table 1.

**Figure 1.**
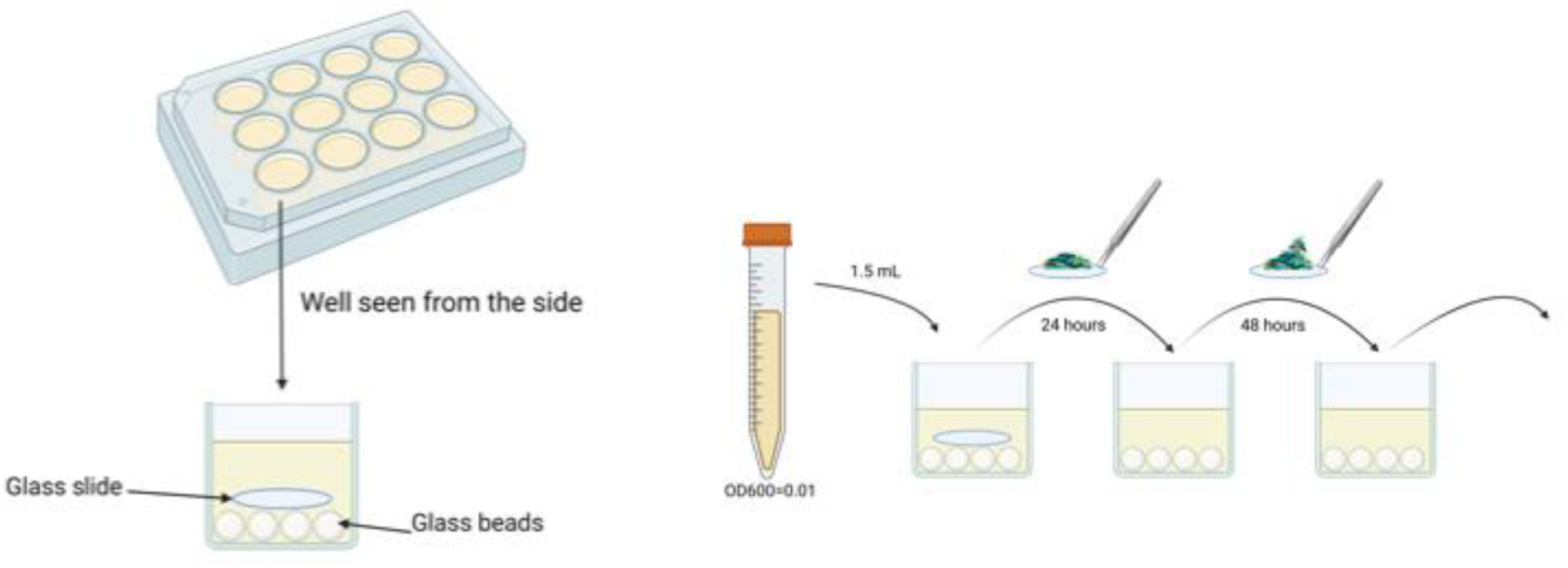
Long-term biofilm cultivation using regular transfers of glass slides to fresh medium. Glass slides acted as surfaces on which bacteria could colonize and form biofilm. To elevate slides from the bottom, a single layer of glass beads (Ø 4 mm) was placed at the bottom of the wells in a 12-well plate. Every 24 h, glass slides were transferred to new wells with fresh medium and in some cases other bacteria or phages.

### Proliferation of phage particles

To proliferate phage stocks, 200µl of the pure phage stock was used to infect a 200ml culture of exponentially growing bacteria (*K. cryocrescens* MB101 or *E. coli* MG1655) in LB supplemented with CaCl_2_ + MgCl_2_ (Final conc. 10 mM). Next day, the culture was centrifuged (5500xG, 4°C, 10 min) and supernatant transferred to a new flask where it was supplemented with NaCl (Final conc. 1.0 M) and PEG8000 (Final conc. 1.6 M). It was allowed to dissolve on a shaking table (100 rpm) for 48 hours at 4°C. The PEG-phage solution was centrifuged (10.000xG, 10 min), the supernatant discarded, and the pellet was dissolved in 6 ml SM buffer (5.8g/L NaCl, 2g/L MgSO_4_•7H_2_O, 0.01% W/V gelatine, 50ml Tris-HCl 1M pH 7.5) by shaking overnight at 4°C to extract purified phage particles. Next day, the solution was centrifuged again and the supernatant filtered (0.22µm pvdf) into sterile glass tubes. Reagents, bacterial strains and phages are listed in Suppl. table 1.

### Image acquisition and analyses

All images were acquired with an inverted LSM800 confocal laser scanning microscope (Carl Zeiss Inc.) equipped with 63X/1.4 oil and 100x/1.4 oil plan-apochromat DIC M27 objectives. For biofilm biovolume quantification, images were acquired with the first objective in an area covering 101.41µm x 101.41µm (1524×1524 px) and a z-interval of 0.7µm. For detailed quantification of local density, we applied the second objective in an area of 63.89µm x 63.89µm (2048×2048 px) and a z-interval of 0.4µm. Image analyses were conducted in the BiofilmQ software [26]. Biofilm biovolume was quantified by thresholding (Otsu’s method) for segmentation and subsequent calculation of default ‘global biofilm properties’. For calculation of local density, we applied a cube-based approach where image segmentations were sliced into a grid of cubes of 93 vox (2.9 µm side length) with Top-Hat filtering at 25 vox for image preprocessing. The cube volume fraction was then used as proxy for cell density.

Co-localisation of *E. coli* (red) and *V. anguillarum* (green) cells was calculated with the parameter function ‘mean shell intensity’ and a custom-made MATLAB script named “Mads_surroundingCells.m” (Supplementary information), which calculates the amount of green cells around every red aggregate of cells in a 15-pixel radius (0.998µm side length). Software details are listed in Suppl. table 1.

### Plasmid construction and genetic manipulations

PCR reactions for cloning procedures were routinely conducted with Phusion Hot Start II polymerase, dNTPs and HF buffer solution according to manufacturer’s description. Verification of mutant construction by colony-PCR was routinely conducted with PCRBIO Taq polymerase and PCRBIO reaction buffer according to manufacturer’s description. For construction of *V. anguillarum* PF4 mutants we used pDM4 as a backbone vector as previously described [27]. The backbone was amplified with primers MFHO170 and MFHO171. Regions flanking the genes of interest were amplified from genomic PF4 DNA with primers designed to overlap and assemble by NEBuilder HiFI DNA assembly (Suppl. table 3). PCR products were routinely purified from gels. All plasmid designs were performed and verified in licensed SnapGene software. All pDM4 derivatives were transformed by electroporation into *E. coli* S17-1 λpir as they are R6K based and requires this gene for replication.

Vector DNA was introduced into *V. anguillarum* by filter mating (0.22um, MCE) conjugation, utilizing the mobilization helper plasmid pRK2013 at ratio 1:1:1 of donor, helper and recipient at 30°C overnight. Cells were washed from the filter in 0.9% NaCl solution and plated on TCBS agar supplemented with 2.5µg/ml chloramphenicol to select for merodiploid transconjugants. Candidate colonies were grown in LB Miller supplemented with 5% sucrose for 3 hours, then diluted 10x and grown for another 2 hours to enrich for the second crossover event before plated on LB Miller agar supplemented with 5% sucrose. To differentiate colonies returning to wildtype genotype from mutants, colony-PCR was performed with primers flanking the gene of interest (Suppl. Table 2). Template DNA was acquired by transferring colony material into 100µl molecular H_2_O and heating at 95°C for 10-15 min. Final verification of mutant candidates was performed by Sanger sequencing (Mix2Seq, Eurofins Genomics) of purified PCR product. Reagents and plasmids are listed in Suppl. table 1.

Construction of mini-Tn7 delivery plasmids for integration of fluorescent markers were also performed by NEBuilder HiFi assembly. Integration of fluorescent markers by mini-Tn7 delivery plasmids were conducted by electroporation of pMFH7 + pTNS2 in *E. coli* AR3110 and by triparental mating with pNUT2703, pTNS2 and pRK2013 in *V. anguillarum* PF4. For the latter, Tn7 positive colonies were selected on TCBS supplemented with 50µg/ml kanamycin. Verification of proper integration was performed by colony PCR and subsequent Sanger sequencing of PCR products with primers MFHO273 and MFHO274 for *V. anguillarum* and MFHO9 and MFHO10 for *E. coli* (Suppl. Table 3). These primers flank the attTn7 site in their respective host. For integration of fluorescent markers by pGRG36 derivate we followed a previously described protocol [28], with a lower final temperature (37°C) for loss of temperature-sensitive plasmid in *K. cryocrescens*. Correct attTn7 integration was verified by colony-PCR and Sanger sequencing. For *E. coli* with primers MFHO9 and MFHO10, and with MFHO69 and MFHO70 for *K. cryocrescens* (Suppl. Table 3).

### Labelling of T7 phage particles

The T7 stock was proliferated by PEG precipitation but with the pellet dissolved in PBS rather than SM buffer, so the Tris components of the buffer would not interfere with labelling. 500ul of the stock was used for labelling with an Alexa Fluor 568 protein labelling kit (Suppl. table 1) following manufacturer’s instructions. The final titer was estimated to be around 5×10^9^ PFU/ml after dialysis.

### Phage adsorption to biofilm communities

Monospecies biofilms were grown for 72h in the glass transfer assay. Wells in a 12-well plate were filled with fresh LM medium and T7 particles (with approx. 4×10^6^ PFU/ml) and mixed thoroughly with the pipette, before biofilm containing glass slides were added. The number of free phage particles were enumerated by plaque assay at four different time points: After 2min, 20min, 2h and 4h. Just before sampling, the plate was shaken on shaking table at 150RPM for even distribution of particles in the medium. 200µl were sampled, centrifuged (6000xG, 4°C, 2min), the supernatant was transferred to a fresh Eppendorf tube and filtered with a syringe filter (0.22µm, PVDF). The filtered sample was diluted in SM buffer and spotted on a lawn of *E. coli* MG1655 in 0.4% topagar.

### Isolation and sequencing of kluyveraphage Lyn

A sewage sample from the wastewater treatment plant Lynetten was centrifuged at 10.000xG for 10 min before supernatant was filtered (0.45µm PVDF) to get rid of larger particles and bacterial cells. 27ml of the cleaned wastewater was supplemented with 3ml of 10X LB, CaCl_2_ + MgCl_2_ (final conc. 10 mM) and 100µl of *K. cryocrescens* MB101 overnight culture was added to enrich for phages infecting this strain and incubated overnight at 37°C. The enrichment was centrifuged, filtered, diluted in SM buffer, and plated using 4 ml 0.4% molten LB top agarose supplemented with CaCl_2_ + MgCl_2_ (Final conc. 10 mM) and 100 µl host cells from an overnight culture on LB plates. Single plaques were picked and transferred into 500µl SM buffer solution, vortexed and left for minimum 1h or overnight at 4°C and then diluted in SM buffer and plated again as described above. This process was repeated three times to ensure a pure phage stock of the phage then named Lyn. To achieve a high titer stock the phage was proliferated and purified by PEG precipitation.

Phage DNA was extracted according to previous descriptions [29,30] by incubating 200µl of high titer stock with 5U/ml of DNase for 1 hour at 37°C. Subsequently 20µl of 50 mM EDTA pH8, SDS (final conc. 0.1%) and 6U of proteinase K was added and the solution was incubated for 1 hour at 55°C. To inactivate enzymes, the solution was heated to 70°C and left for 10 min before the DNA was purified with the DNA Clean and Concentrater-5^TM^ kit. Sequencing library was built using NebNext FS II and the library was sequenced on the Illumina iSeq100 platform (150 base pair, paired end). The genome was assembled using both CLC Genomics Workbench and SPAdes. Phage Lyn was predicted to have direct terminal repeats (DTR), identified by spikes in read start coverage, as previously explained elsewhere [31] and the genome start position was set accordingly. The 39408 bp genome was submitted to taxMyPhage to identify closest relative and submitted to pharokka for visualization and annotation of the genome. Reagents and software details are listed in Suppl. table 1.

### Sequencing of bacterial strains

Although the genome sequence of *V. anguillarum* PF4 was already publicly available (Suppl. table 2), we re-sequenced our isolate to ensure that we were working with the correct strain and that it did not encode *rbmAB*. This resulting sequence was identical to the publicly available and verified lack of *rbmAB* in the strain used in this study. DNA from overnight cultures of *K. cryocrescens* MB101 and *V. anguillarum* PF4 was extracted using DNeasy Ultraclean Microbial kit according to manufacturer’s instructions. Genomes were sequenced with Illumina MiSeq platform by sequencing Nextera XT libraries obtained from each isolate. Quality control and trimming was conducted in FastP. Assembly of reads for the *V. anguillarum* PF4 genome was conducted using Unicycler with a mean coverage of 132x. The genomic DNA of *K. cryocrescens* MB101 was additionally sequenced with the Nanopore long-read platform to obtain a fully closed genome. The library was prepared with rapid barcoding kit 96 and sequenced as part of the PromethION flowcell FLO-PRO002 (based on R9.4.1 pores), following the manufacturer’s instructions. Data was basecalled with Guppy with “super-accurate” setting. Filtlong was used for filtering, removing the worst 5% of reads and reads shorter than 1000bp. Hybrid genome assembly was performed in Flye with a mean coverage of 130x. Genome quality was verified with CheckM, with no sign of contamination. The taxonomy of MB101 was analysed using the TYGS genome type server (GBDP whole-genome-based) and thereby identified as *K. cryocrescens*. Genomes have been deposited to NCBI under BioProject ID PRJNA1266407. *V. anguillarum* PF4 as GenBank accession JBOCIA000000000, with *vps* clusters on contig 3. Reagents and software details are listed in Suppl. table 1.

### Attesting the ability to form curli with optotracers

EbbaBiolight680 optotracer was added to LB w.o. salt (1:500 ratio) supplemented with 1.4% noble agar and 2ml was placed in wells of a 6 well plate. After agar was solidified, 10ul of bacterial overnight culture was spotted to the centre of the well. Fluorescence was monitored in a Synergy microplate reader (Biotek) with fluorescence signal acquired every second hour (excitation peak λ 561 nm, emission peak λ 680 nm).

### Statistical analyses

All statistical tests were performed in GraphPad Prism. For data in Fig. 2 we compared phage-exposed to buffer exposed with the application of a two-way ANOVA with Fisher’s LSD test for comparison. For the remaining data, we performed Brown-Forsythe and Welch ANOVA, which do not assume equal standard deviation, and performed Dunnett’s T3 multiple comparison test for respective comparisons. For the shielded fraction, the Kruskal-Wallis test was applied as data was not normally distributed, followed by Dunn’s test for multiple comparison.

**Figure 2.**
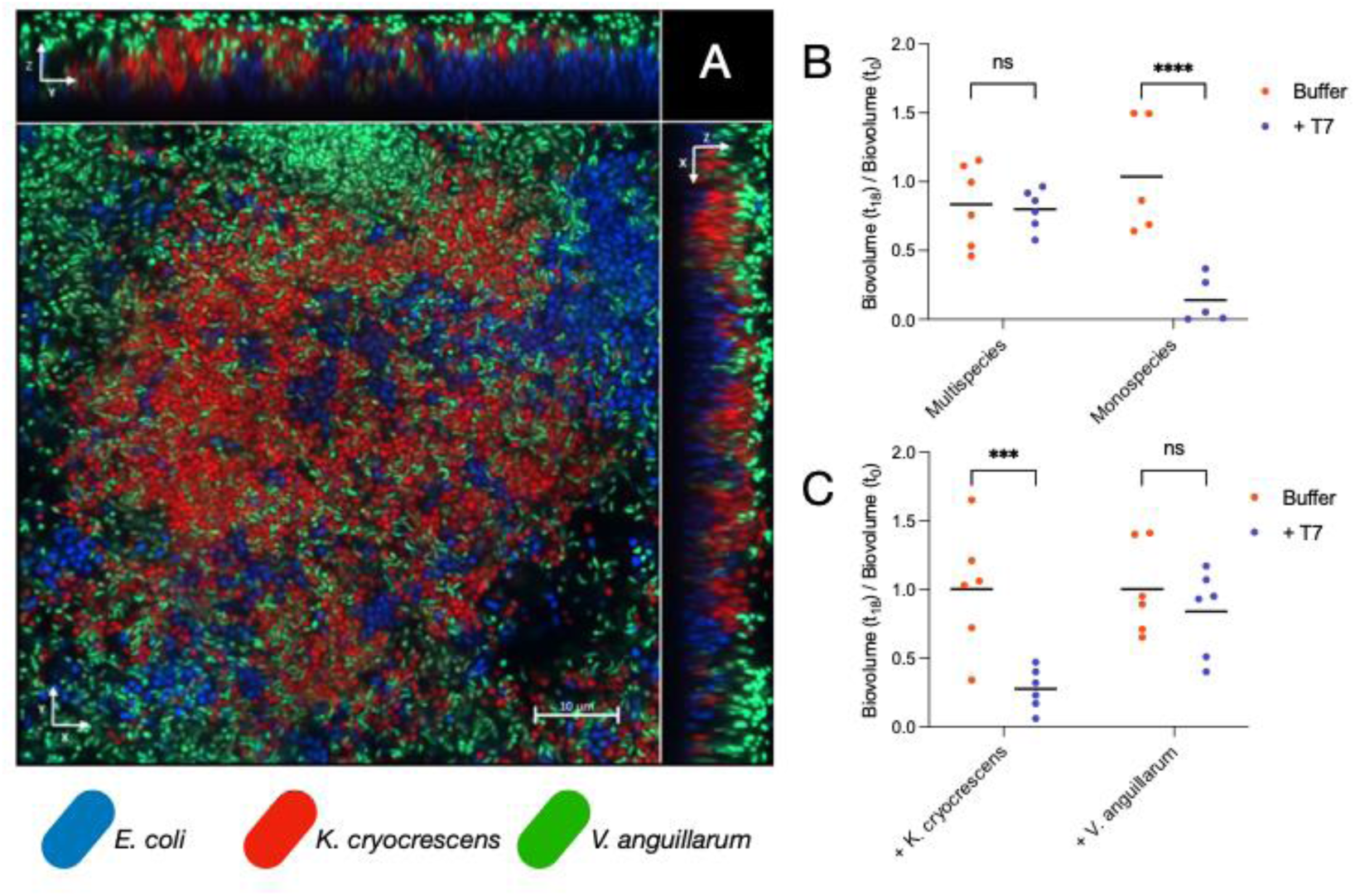
Multispecies biofilm community members were protected from phage infection. **A)** Orthogonal view of a three-species biofilm comprised of *E. coli* (blue), *K. cryocrescens* (red), *V. anguillarum* (green) where the central window represents a cross section in the middle of the community. The scale bar represents 10 µm. **B)** The biovolume of *E. coli* was unaffected by 18 hours of exposure to the lytic coliphage T7 when grown in the multispecies community, whereas *E. coli* was almost eradicated when growing in a monospecies biofilm. **C)** Dual species biofilms of *E. coli* with *K. cryocrescens* or *V. anguillarum*, respectively, revealed that *V. anguillarum* was able to protect *E. coli* from phage infection, while *K. cryocrescens* was unable to protect *E. coli*, indicating that *V. anguillarum* facilitated the protection in the multispecies community (ANOVA, Fisher’s LSD test for comparison, P<0.001 = ***, P<0.0001 = ****).

## Results

### Establishing a 3-species biofilm model

To examine the effect of phages in multispecies biofilms, it was essential to have a multispecies community that could be consistently established. As a model system, we modified the bead-transfer assay originally developed by Poltak & Cooper [32], because this system enables continuous biofilm cultivation and is not restricted in time. Instead of the glass beads that were originally used by Poltak & Cooper [32], we used glass slides, because their flat architecture enables easy analysis of spatial biofilm organization by microscopy. Furthermore, instead of replacing glass beads for every transfer, we transferred the same glass slides from well to well throughout the entire incubation period, to minimize rounds of dispersal and subsequent re-attachment. Others have previously used a similar approach to study the phage susceptibility of biofilms up to twenty days of age [33]. To elevate the glass slide, a single layer of glass beads was placed at the bottom of the wells (Fig. 1).

The bacterial community consisted of *Vibrio anguillarum* PF4, *Escherichia coli* AR3110, and *Kluyvera cryocrescens* MB101. These three strains are all biofilm formers and were genetically manipulated to encode distinct fluorescent reporters on the chromosome. Initially, *V. anguillarum*, *E. coli,* and *K. cryocrescens* were mixed and inoculated simultaneously. However, this led to the formation of biofilms that were heavily dominated by *V. anguillarum* with only a few randomly distributed aggregates of *K. cryocrescens* and *E. coli* (Suppl. Fig. 1). Instead, when species were added in sequential order (also known as priority affects) [34], first letting *E. coli* colonize the glass for 24 hours, then adding *K. cryocrescens* and finally introducing *V. anguillarum,* the three species were all present in the biofilm. The order in which species were added was also reflected in the resulting community structure, with *E. coli* at the bottom, *K. cryocrescens* in the middle parts and *V. anguillarum* at the top (Fig. 2A, Suppl. Fig. 2).

### Multispecies community protected against phage infections

To test how multispecies community properties impacted the dynamics of phage infection, coliphage T7 was added to monoculture *E. coli* biofilms and to the multispecies model community. The T7 phage only infected the *E. coli* cells (Suppl. Fig. 3), and the biomass of *E. coli* was dramatically reduced in monospecies biofilms after 18 hours of exposure to T7 (Fig. 2B). However, when growing in the presence of the two other species in the multispecies community, the *E. coli* biomass was unaffected by the presence of T7 (Fig. 2B). To elucidate whether the presence of both species was required for protection or whether a single biofilm-forming neighbour was sufficient, *E. coli* was co-cultured with only *V. anguillarum* or *K. cryocrescens*, respectively. In such dual-species *E. coli-K. cryocrescens* biofilms, *K. cryocrescens* did not provide any phage protection, and *E. coli* cells were killed by the T7 phage (Fig. 2C) similar to *E. coli* monoculture biofilms. In contrast, in phage exposed dual-species *E. coli*-*V. anguillarum* biofilms, *E. coli* was protected and the biovolume reduction was not greater than in the control where SM buffer was added instead of phages (Fig. 2C).

### Phage protection required all biofilm matrix genes

The reduction of biomass in phage-exposed monospecies *E. coli* biofilms was greater than expected with almost complete eradication in some replicates (Fig. 2B). Previous studies have described how biofilms of *E. coli* AR3110 older than 48-60 hours produce a matrix containing curli amyloid fibres that protect cells and sequester phages at the periphery of the community [18,19,35]. As the *E. coli* cells in our experiments had been incubated for a total of 72h before phage exposure, this should have been sufficient time to produce the protective curli. The use of curli-specific optotracers verified that the fluorescently labelled strain used here was able to produce curli (Supplementary fig. 4). Furthermore, the expression of the *csgBAC* operon was confirmed to be temperature-responsive [36,37] (Suppl. Fig 5A Suppl. Fig. 6), and immunostaining of CsgA-His revealed that the curli fibres, although being expressed at a transcriptional level, did not assemble in the matrix in NaCl containing medium (Suppl. Fig. 5B-D). Thus, we conclude that in our glass slide transfer assay, conducted in LB medium with 1% NaCl at 30 °C (vs. no salts and room temp. in previous studies [18,19,35]), killing of *E. coli* killing by T7 phages in monospecies communities was possible due to absence or low abundance of curli amyloid fibres in the biofilm matrix.

Next, we sought to investigate the mechanism behind the observed phage protection of other species in the community conferred by *V. anguillarum*. Matrix components, including amyloid fibres [18] and exopolysaccharides [17], can protect against phage infection in monospecies biofilms. Hence, we hypothesized that the phage protection conferred by *V. anguillarum* was due to its production of one or several specific matrix components. Interestingly, our strain of *V. anguillarum* did not encode *rbmA* (Suppl. Fig. 9), previously found to ensure a dense packaging of cells in *V. cholerae* biofilms [38,39] and essential for *V. cholerae*-facilitated T7 protection of *E. coli* [19]. A comparative bioinformatic analysis of the two *vps* (Vibrio polysaccharide) gene clusters, and the region between these, in which the *rbmABC* genes are located in *V. cholerae* [40], revealed that most *V. anguillarum* strains lack *rbmA* and *rbmB* but encode *rbmC*: Only one out of thirteen strains encoded *rbmAB* (Suppl. Fig. 9). This led us to conclude that other matrix components, rather than RbmA, conferred the phage protection of *E. coli* by the *V. anguillarum* strain used in this study. To test if a specific matrix component was responsible for phage protection, we constructed an array of *V. anguillarum* matrix-mutants by in-frame deletion, each mutant lacking a matrix-associated gene or cluster. Firstly, we quantified the ability of each mutant to form biofilms (Fig. 3A). Here, the *bap1*-, *vpsL*-, and *vpsIJ*-mutants were unable to form mature biofilms, suggesting that these genes are essential for biofilm formation. In contrast, mutants lacking *rbmC*, the *vpsMNOP*-cluster or the putative *csgD* homologue formed biofilm in volumes similar to wildtype (Fig. 3A). Interestingly, Bap1 and RbmC have redundant functions in *V. cholerae* biofilms, and knockout of *bap1* only reduces biofilm formation slightly in *V. cholerae*, while it requires a double knockout mutant to have a biofilm deficient *V. cholerae* strain [41]. As the single *bap1* deletion was sufficient to disable biofilm formation of *V. anguillarum*, we speculated whether *rbmC* in *V. anguillarum* could be mutated and non-functional. Alignment to the nucleotide sequence of *V. cholerae* N16961 *rbmC* (VC0930) showed 297 mismatches, but no deletions, gaps or nonsense mutations, and on the amino acid level the RbmC proteins were 94% identical (Suppl. Fig. 10A). On the other hand, Bap1 of *V. anguillarum* aligned poorly with *V. cholerae* Bap1 (VC1888) and was only 69% identical on the amino acid level (Suppl. Fig. 10B).

**Figure 3.**
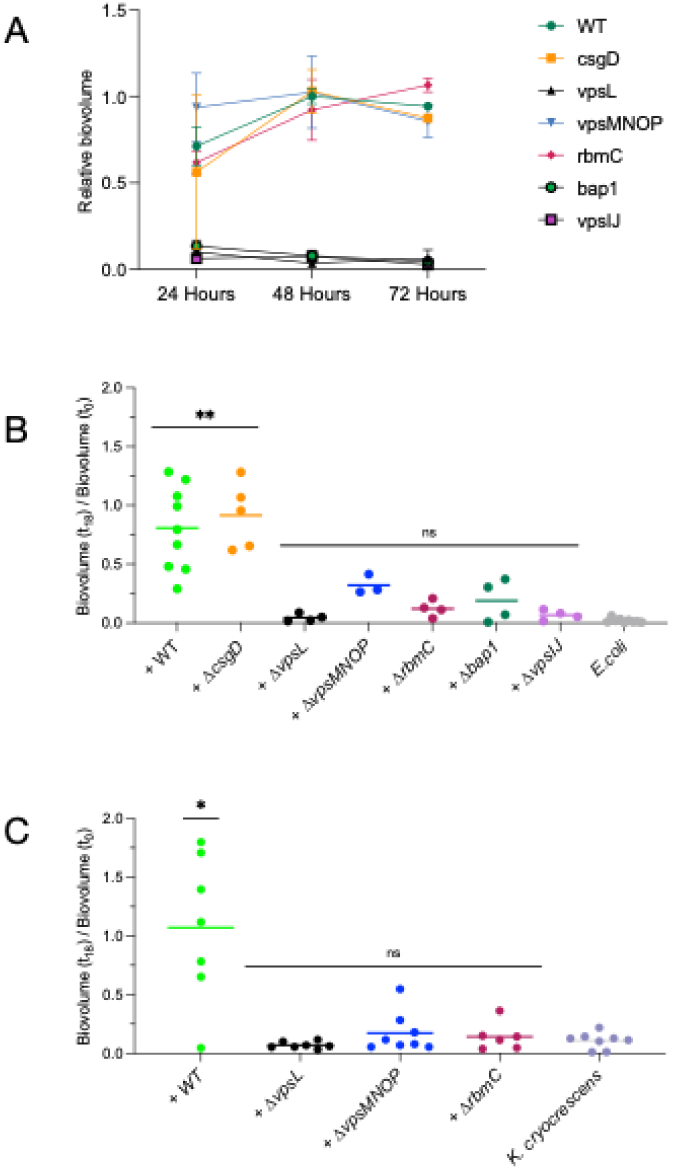
Biofilm formation of *V. anguillarum* mutants and their ability to protect from phage infection. **A)** The temporal biofilm formation was quantified in monocultures of mutants of *V. anguillarum*, where deletion of *vpsL*, *vpsIJ* and *bap1* matrix-encoding genes made this species unable to form a biofilm, while mutants with deletion of matrix-encoding genes *vpsMNOP* and *rbmC* and the putative transcription factor csgD produced biofilm volumes similar to the wildtype (Error bars represent S.E.M.). **B)** *V. anguillarum* wildtype and Δ*csgD* were able to provide a significant level of protection of *E. coli* during phage exposure, while the Δ*vpsL*, Δ*vpsMNOP*, Δ*rbmC*, Δ*bap1* and Δ*vpsIJ* mutants did not yield significant protection compared to *E. coli* growing in monoculture. **C)** The same pattern was observed when replacing *E. coli* with *K. cryocrescens*, where a subset of *V. anguillarum* mutants (Δ*vpsL*, Δ*vpsMNOP*, Δ*rbmC*) were unable to protect *K. cryocrescens* from infection and killing, while the wildtype provided significant protection compared to the *K. cryocrescens* monoculture (ANOVA, Dunnett’s multiple comparison with the respective monoculture as reference. P<0.05 = *, P<0.01 = **).

As expected, the biofilm-deficient mutants (Δ*bap1*, Δ*vpsL*, Δ*vpsIJ*) were unable to protect *E. coli* from phage T7 infection and the biomass was reduced to levels similar to *E. coli* monoculture biofilms infected by phage T7 (Fig. 3B). Remarkably, two of the mutants that were still able to form biofilms (Δ*rbmC* and Δ*vpsMNOP*) did not protect *E. coli* from phage infection, meaning that only the Δ*csgD* mutant and wildtype protected *E. coli* (Fig. 3B). CsgD is not a matrix component per se, but a transcription factor known to regulate matrix production in *E. coli* [42–44] and *Salmonella typhimurium* [45], where it orchestrates the expression of amyloid fibres among other components[46]. Since *V. anguillarum* did not produce amyloid fibres (Suppl. Fig. 4A), and the fact that this gene does not impact biofilm volume or interspecies phage protection could indicate that this putative gene is not biofilm-related in *V. anguillarum*.

Analogous to the lack of phage T7 protection provided by the matrix mutants of *V. anguillarum* to *E. coli*, these *V. anguillarum* mutants also did not protect *K. cryocrescens* from phage infection, while the *V. anguillarum* WT secured a significant level of *K. cryocrescens* survival (Fig. 3C). The phage used in this experiment was isolated from the wastewater treatment plant Lynetten, where wastewater from Copenhagen and surrounding municipalities is treated, and hence this phage was given the name Lyn. Bioinformatic analysis suggested that kluyveraphage Lyn was part of the Autographiviridae family and the Foetvirus genus, with the closest relative being coliphage SRT7 (Accession no. MH370477) to which it shared 94.3% similarity on nucleotide level. The phage is further characterized in Suppl. Fig. 11.

### Phage protection by high cell density and tight co-localization

Since *rbmC* and *vpsMNOP* were essential for phage protection of *E. coli* and *K. cryocrescens,* but not for biofilm formation, we hypothesized that these genes facilitated the ability to confine and trap phages in the *V. anguillarum* matrix. To test this hypothesis, we labelled the T7 phage particles by fluorophore conjugation and visualized phage particles in mono- and dual-species biofilms. However, the T7 phage particles did not attach to or entangled in the *V. anguillarum* matrix. In contrast, we observed a strong signal of T7 particles adsorbed to the monoculture biofilms of *E. coli* (Suppl. Fig. 7). This suggests that the *V. anguillarum* matrix functions as a barrier, rather than trapping aggregates of phages in the matrix. This observation was supported by a phage adsorption assay, where we quantified the number of free T7 particles over time in wells with *V. anguillarum* biofilm formed on glass slides. Here, the numbers of free T7 particles in the medium were similar to those in the control with sterile glass slides (Suppl. Fig. 8), indicating that phage particles were not trapped in the matrix.

Inspired by the previously reported importance of tight cell packaging of *V. cholerae* biofilms in the protection of *E. coli* [19], we quantified the cell density of *V. anguillarum* WT, Δ*vpsMNOP* and Δ*rbmC* biofilms, respectively. For this analysis, images were cube-segmented using BiofilmQ [26] and the average fraction of the cube volume occupied by cellular biovolume was calculated as a proxy of cell density. The wildtype biofilm was significantly denser than the biofilm of the two mutants (Fig. 4A-D), which might explain why the mutants did not protect *E. coli* and *K. cryocrescens* from phage exposure. To interpret the impact of density at a local level, the ability to surround and shield individual clusters of *E. coli* cells was quantified. Specifically, a 1 µm shell surrounding the red cells (*E. coli*) was computed, and the fraction of the shell filled with green cells (*V. anguillarum*) was calculated and termed the shield fraction (Fig. 4E). This analysis revealed that clusters of *E. coli* cells were significantly more surrounded in their vicinity by wildtype *V. anguillarum* cells than when co-cultured with one of the mutants (Δ*vpsMNOP* and Δ*rbmC*) (Fig. 4F). This could potentially explain why T7 was prevented from accessing *E. coli* cells when *V. anguillarum* formed intact biofilms on top. Another aspect of this spatial community dynamic is that we repeatedly observed that *E. coli* translocated upwards in the z-direction in unfilled voids in the community of the mutants (Fig. 5G-H). This interaction was rarely observed in the presence of the *V. anguillarum* WT, and never to the same extent. This may indicate that the lower density of the biofilms formed by the mutants enabled *E. coli* to grow, sometimes also on top of *V. anguillarum*, which created areas accessible for T7 adsorption and subsequent access the lower parts of the biofilm, which would otherwise be shielded from exposure.

**Figure 4.**
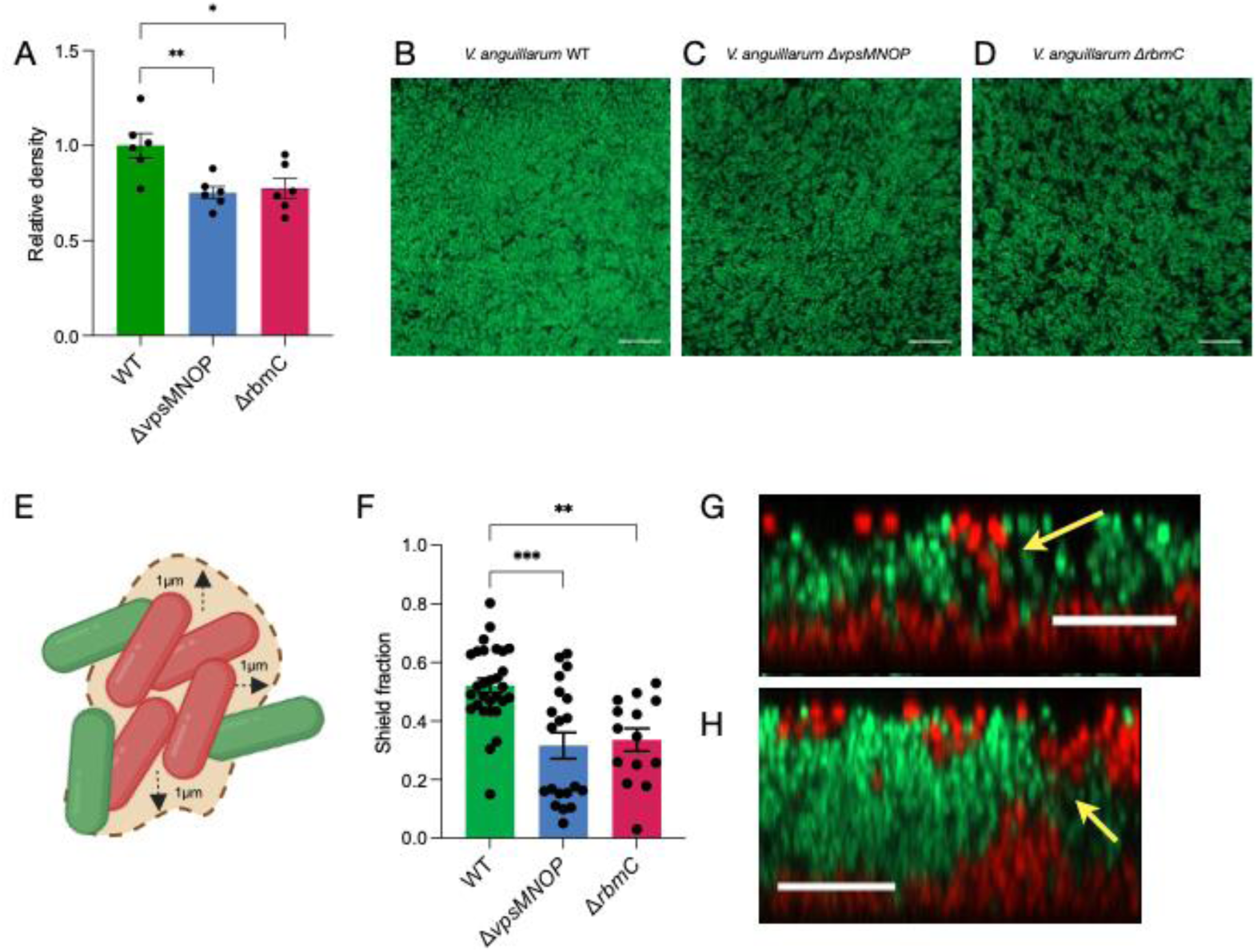
Community architecture changed in biofilms formed by *V. anguillarum* matrix mutants. **A)** The biofilm cell density was lower in the two mutants (Δ*vpsMNOP* and Δ*rbmC*) than in the wildtype (measured in monospecies biofilm) (ANOVA, Dunnett’s multiple comparison. P<0.05 = *, P<0.01 = **). **B-D)** Representative images of biofilm formation in the XY planes of *V. anguillarum* WT (B), Δ*vpsMNOP* (C) and Δ*rbmC* (D), respectively. Scale bars represent 10µm. **E)** Schematic illustration of the concept of shield fraction and how it was calculated around red cells. **F)** *E. coli* cells growing with the wildtype *V. anguillarum* were shielded more than *E. coli* cells growing with the *V. anguillarum* Δ*vpsMNOP* or Δ*rbmC* mutants, respectively (Kruskal-Wallis, Dunn’s multiple comparison. P<0.01 = **, P<0.001 = ***). **G-H)** Red *E. coli* cells were found to translocate in the Z direction, growing upwards in voids of the *V. anguillarum* matrix (yellow arrow) of the Δ*vpsMNOP* (G) and Δ*rbmC* (H) mutants, enabling growth on top of *V. anguillarum* cells (green) and might create an entry for phage infection in the community. Scale bars represent 10µm.

**Figure 5.**
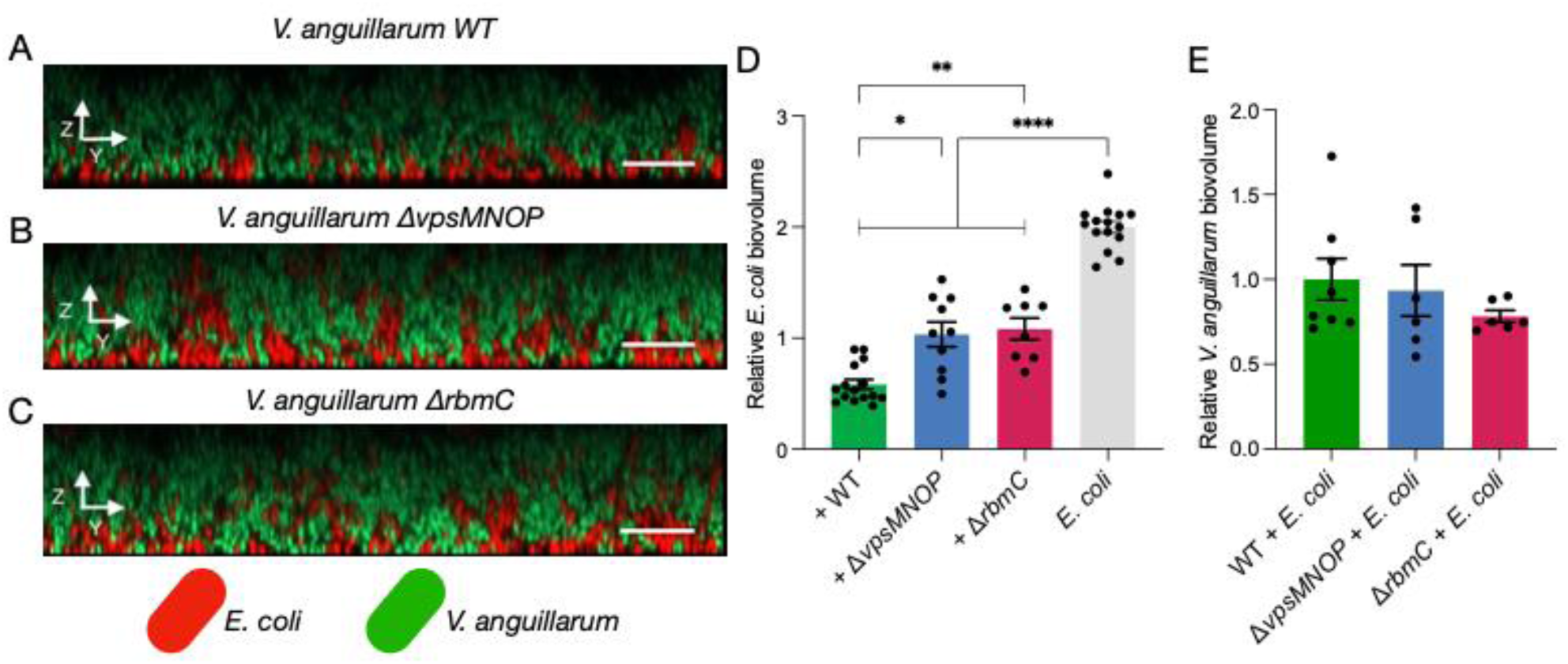
Phage protection comes with a competitive cost. **A-C)** Representative images of *E. coli* cells (red) in biofilm community with *V. anguillarum* (green) WT cells (A), Δ*vpsMNOP* cells (B) and Δ*rbmC* cells (C). Scale bars represent 10 µm. **D)** The relative biovolume of *E. coli* was reduced when co-cultured with *V. anguillarum,* and even further reduced when growing with the wildtype, compared to when growing with one of the two mutants (Δ*vpsMNOP* and Δ*rbmC*) (ANOVA, Dunnett’s multiple comparison. P<0.05 = *, P<0.01 = **, P<0.0001 = ****). **E)** The biovolume of *V. anguillarum* mutants was unaffected by co-cultivation with *E. coli* as the relative biovolume of the mutants was similar to the wildtype.

If *V. anguillarum* WT cells hindered the expansion of *E. coli* in the z-direction, while the Δ*vpsMNOP* and Δ*rbmC* mutants did not, the abundance of *E. coli* in the biofilm should be greater when incubated with one of the mutants compared to the wildtype. This hypothesis was supported by comparing the biomass of *E. coli* across the three scenarios. Co-cultivation reduced *E. coli* abundance, with a further reduction observed when incubated with the wildtype compared to the less dense mutants (Fig. 5D). Hence, *V. anguillarum*-mediated protection of *E. coli* from phage exposure is associated with a cost of *E. coli* cell growth. Thus, protection was likely facilitated by spatial constraints by two means; 1) The dense cell packaging prevented phage access to *E. coli* cells, and 2) *E. coli* cells were unable to grow into areas exposed to phages. To comprehend the competitive species dynamics, we also compared the abundance of *V. anguillarum* mutants when co-cultivated with *E. coli*. Here, there was no significant difference in abundance of wildtype and mutants indicating that the deletion of matrix component encoding genes did not alter *V. anguillarum* growth in the dual species community (Fig. 5E). In conclusion, the tight cell packaging of the *V. anguillarum* WT was necessary to hinder *E. coli* from expanding its population, while the less dense biofilms formed by mutants (Δ*vpsMNOP* and Δ*rbmC*), despite reaching abundances similar to the wildtype, enabled *E. coli* expansion in the less compact pockets of the mutant communities.

## Discussion

Traditionally, bacteria-phage dynamics have been studied in simple contexts with a single bacterial species exposed to phage particles or in complex metagenomic studies with low resolution [47]. To include community context in the interplay of bacteria and phages, the dynamics in a multispecies biofilm community were studied here. In a study by Winans et al. [19], tight cell packaging of *V. cholerae* was a key determinant for precluding phage infection. Here, this architectural nature was attributed to *V. anguillarum*, which protected *E. coli* and *K. cryocrescens* from infectious killing by forming a dense biofilm. Interestingly, this study highlighted significant differences between the genes involved in formation of a biofilm matrix of *V. anguillarum* and *V. cholerae*. First of all, most *V. anguillarum* strains, including the strain used here, lack the gene *rbmA* (Suppl. Fig. 9), which encodes the scaffolding protein that binds VPS and facilitates tight assembly of cells in the matrix [39], and is required for *V. cholerae*-facilitated protection of *E. coli* [19]. Next, the Bap1 and RbmC matrix proteins perform redundant functions in *V. cholerae*, and both need to be deleted in order to observe a strong phenotypic change in biofilm formation [41,48]. Here, Bap1 was found to be an essential matrix component of *V. anguillarum* on its own (Fig. 3A) and shared limited similarity with *V. cholerae* Bap1 on an amino acid level (69% - Suppl. Fig. 10). As *rbmC* was not conspicuously defective (Suppl. Fig. 10), it could suggest that the *V. anguillarum* Bap1 protein comprises other biofilm-essential functions than what has been described for *V. cholerae*. Finally, genes encoding VpsM, VpsN and VpsO are all essential for VPS synthesis and biofilm formation in *V. cholerae* [49]. In this study, the *V. anguillarum* mutant lacking all three genes and *vpsP*, still formed biofilm (Fig. 3A, Fig. 5E). All in all, there seems to be fundamental differences in how respective matrix-associated genes impact the formation of biofilm in the two species.

In this study, it was not possible to achieve consistent and comparable biofilm communities without applying priority effects. As described, simultaneous inoculation produced variable communities, with only a few aggregates of the two other species embedded within a *V. anguillarum* community. The spatial organization changed in biofilms comprising the *V. anguillarum* Δ*vpsMNOP* and Δ*rbmC* mutants, respectively, resulting in *E. coli* exposure to phages in the top layers of the biofilm (Fig. 4G-H). The observation of *E. coli* growing in the less dense voids when co-cultivated with a matrix mutant (Fig. 4G-H), also explains the bimodal distribution of shield fraction for the mutant strains, especially the Δ*vpsMNOP* mutant (Fig. 4F). The cells present at the top are naturally not as surrounded by *V. anguillarum* as those present within or at the bottom of the community. Since the phage protection was lost with this spatial change, it is likely that interspecies shielding in general depends on the order and timing of species’ arrival. One could also argue that sequential arrival of species likely represents natural conditions to a higher degree than simultaneous inoculation. The formation of dental plaques is one example of community assembly in nature. Here, early colonizers foster an environment to which others colonize, increasing diversity over time with a non-random distribution [50–52]. This highlights the need to consider priority effects for community assembly studies [34] and also when studying bacteria-phage interactions.

The formation of curli in the matrix of monospecies *E. coli* biofilm protects against phage infection [18,19,35]. However, since expression and folding of curli is affected by temperature and salt levels, there are many environments, such as the gut of warm-blooded animals, where *E. coli* will be vulnerable to phage exposure, and hence rely on protection provided by other community members. The model system used here has contributed to a broader understanding of the mechanisms at play in bacteria-phage dynamics and interspecies competition in a biofilm context. As this study emphasizes, there is a difference in the community-intrinsic properties of a biofilm community depending on the species present, i.e. co-cultivation with *K. cryocrescens* did not yield any protection. Future work could benefit from including frequent biofilm co-inhabitants from relevant environments to reveal the full impact of interspecies interactions in the interplay with phages and increase validity for potential applications such as microbiota manipulation or clinical phage therapy. One model of interest could be wound infections, which are frequently polymicrobial [22,51]. The next steps in this line of clinical research could benefit from an increase in complexity and introduce organoids or animal models to also include the impact of eucaryotic host cells and immune system [53,54]. Our work adds to an increasing acknowledgement of the importance of community-intrinsic properties and community-context [5,6,8,47] in a perspective of phage-bacteria interaction dynamics. The fact that *V. anguillarum* was able to protect *E. coli* from phage infection, while *K. cryocrescens* was not, and deleting matrix-related genes tampered with this protection, emphasizes that bacteria-phage dynamics depend on the species present, how they interact in the community and the biofilm architecture.

## Acknowledgement

We would like to thank Jesper Juel Mauritzen for helping identifying matrix-associated genes in *V. anguillarum*. We also thank Prof. Mathias Middelboe for sharing *Vibrio anguillarum* PF4. We would like to thank Prof. Dr. Regina Hengge for sharing *E. coli* AR3110 and Dr. Ronja Offer for taking care of the practical aspects of shipping this strain. We would like to thank Prof. Sylvain Moineau and lab manager Denise Trembley from the Felix d’Herelle reference centre in Canada for their effort in maintaining a collection of bacteriophages and sending the T7 coliphage.

## Author contributions

M.F.H. and M.B. conceived the project, M.F.H., J.H.O., A.P.N. and S.L.T. conducted experiments and acquired CLSM images, A.M.D and N.R.H. isolated and characterized Kluyveraphage Lyn, A.M.D., J.H.O. and M.F.H. performed bioinformatics analyses, M.F.H., J.H.O., H.J. and E.J.S analysed and quantified CLSM images, H.J. wrote the customized script for shield fraction quantification, and M.B., L.H.H., K.D. and C.N. supervised the project. M.F.H., M.B. and J.H.O. wrote the original draft. All authors contributed critically to the draft with corrections.

## Data availability

The customized MATLAB scripts for quantification of shield fraction can be found as supplementary information. Specific data reported in this paper will be shared upon request to corresponding author, Mette Burmølle.

