## Supplementary Material for "Biofilm compactness and spatial community composition determine bacterial phage-susceptibility"

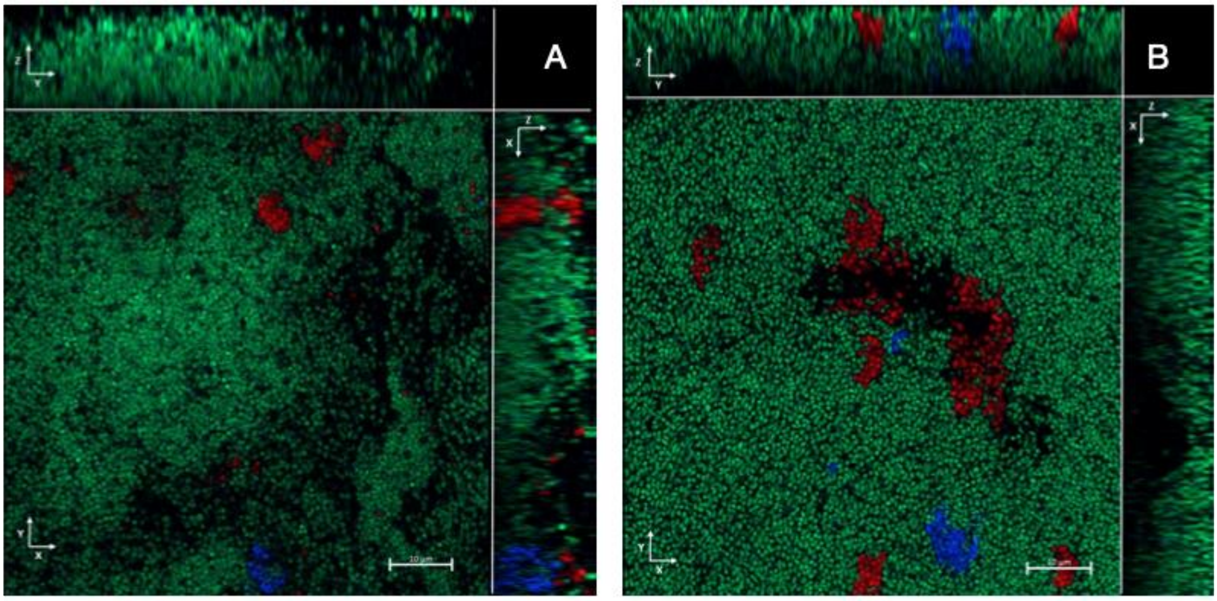


**Supplementary Figure 1) Simultaneous inoculation yields *V. anguillarum* domination.** When all three strains were inoculated simultaneously, the community was dominated by *Vibrio anguillarum* (green), with only few, randomly distributed aggregates of *Kluyvera cryocrescens* (red) and *Escherichia coli* (blue). **AB)** Representative images of two different biofilm communities where the three strains were inoculated simultaneously. Images were acquired 72h after inoculation.

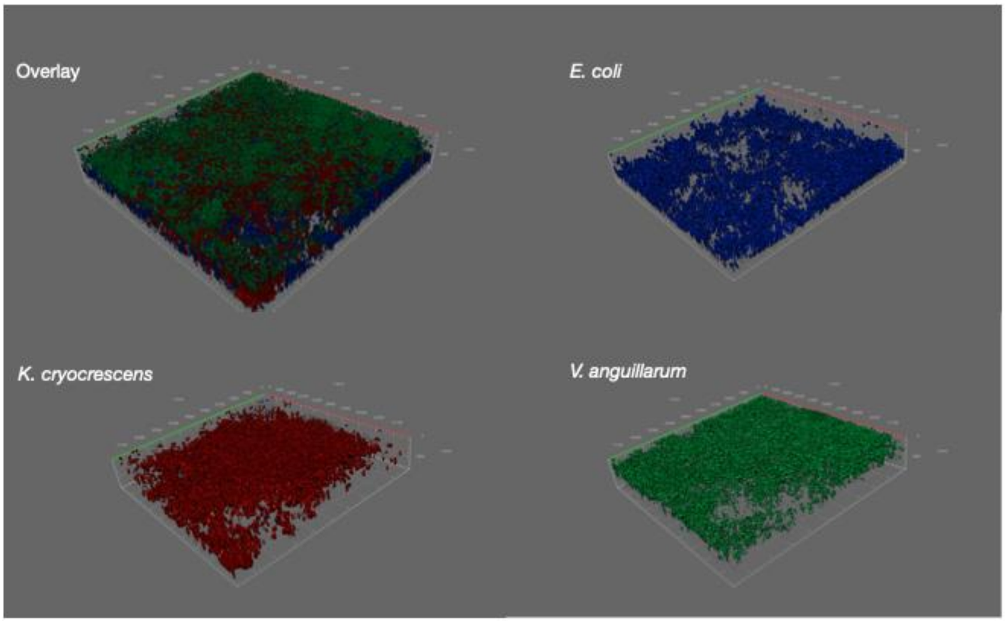


**Supplementary Figure 2) Three-dimensional representation of multispecies community.** The biofilm community of *E. coli* (mTagBFP2/blue), *K. cryocrescens* (mCherry/red) and *V. anguillarum* (sfGFP/green) represented in combination and as single channels, respectively. The biofilm community was established by sequential inoculation, where *E. coli* incubated as monoculture for 24h before *K. cryocrescens* was added and then incubated for another 24h before *V. anguillarum* was added. The order of species addition is reflected in the spatial community composition with *E. coli* mainly distributed in the bottom, *K. cryocrescens* in the middle and then *V. anguillarum* on top. This image was acquired with airyscan detection mode for increased resolution for improved 3D demonstration.

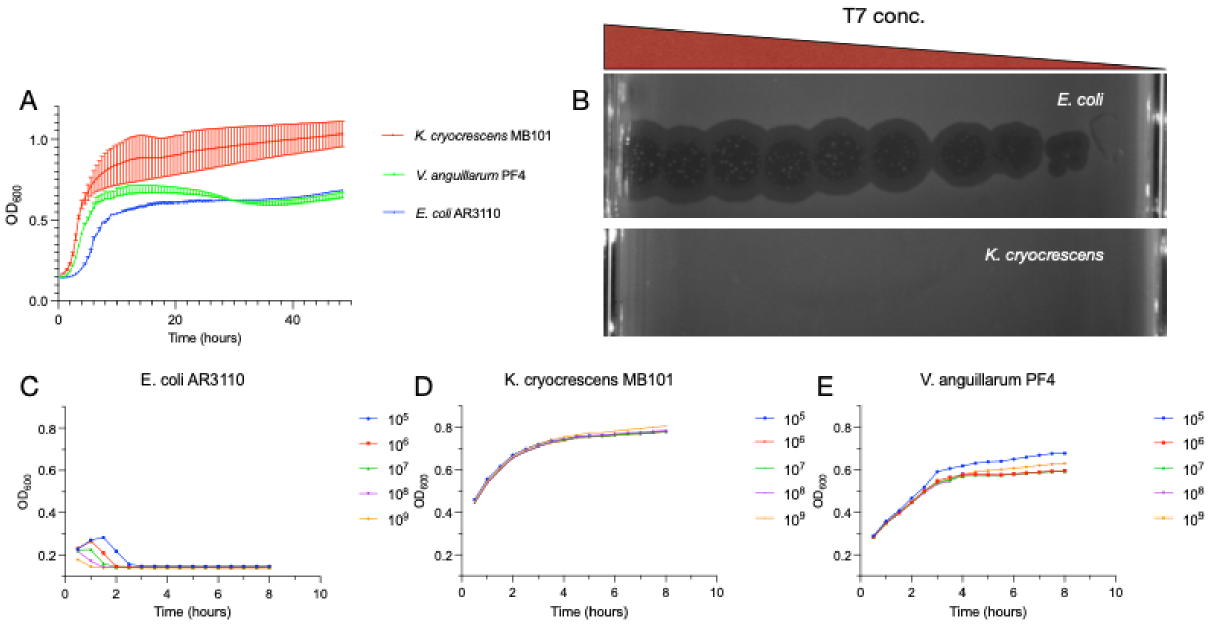


**Supplementary Figure 3) Specificity and infection kinetics of coliphage T7. A)** Growth kinetics of the three strains used in this study in planktonic culture**. B)** Plaque formation assay at decreasing concentration of coliphage T7. Plaque forming units were only observed on lawns of *E. coli.* **C)** Temporal growth kinetics of *E. coli* exposed to coliphage T7 at different concentrations. As the concentration of the lytic phage increases, the population killing is accelerated. **D)** Temporal growth kinetics of *K. cryocrescens* exposed to coliphage T7 at different concentrations. The growth is unaffected by the presence of T7. **E)** Temporal growth kinetics of *V. anguillarum* exposed to coliphage T7 at different concentrations. There is some variability in growth, but the strain continues to grow in all concentrations of T7 tested. Graphs represent the average of three replicates, with error bars representing S.E.M.

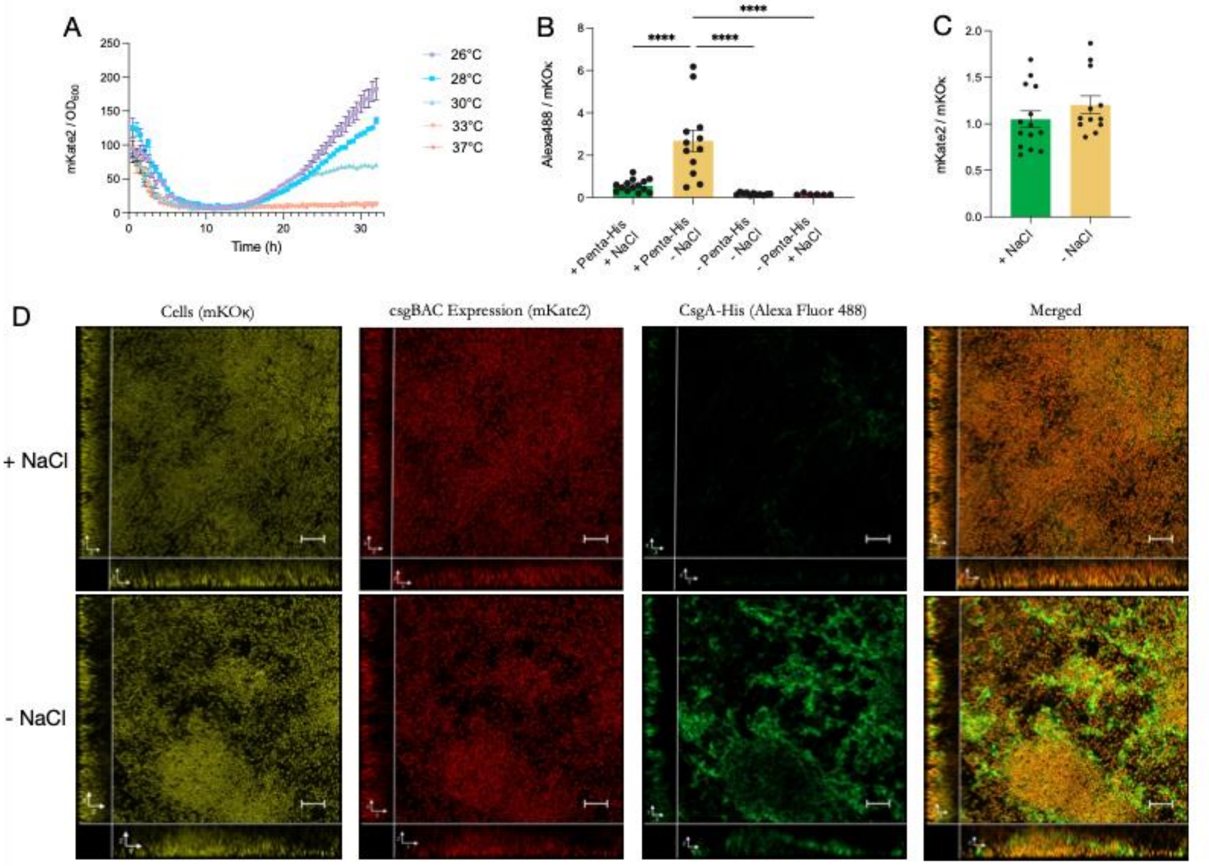


**Supplementary Figure 4) Curli production depends on temperature and NaCl presence. A)** Temporal monitoring of *csgBAC* transcription (mKATE2) at different temperatures, with expression normalized to account for bacterial density ($OD_{600}$). The expression of the operon was temperature sensitive and highest at lower temperatures (26°C) and declined with elevated temperature until absent from 33°C and up. Non-normalized data can be found in suppl. fig. 6. **B)** Quantification of CsgA-His (*csgA*-6xHis) deposited in the matrix by anti-His antibody labelling (Alexa Fluor 488) normalized to the total biomass present (mKOκ) revealed that curli was not present in *E. coli* biofilms grown on glass slides at 30°C in LB medium. Removing salt from the medium allowed for amyloid fibres to form in the matrix, indicating that salt is the determining factor of curli formation (error bars represent S.E.M.) (ANOVA, Dunnett's multiple comparison test, $P<0.0001$ = ****). **C)** Transcriptional expression levels of *csgBAC* (mKATE2) were similar in *E. coli* biofilms in LB medium and LB medium without salt, respectively, indicating that the difference between the two scenarios was due to post-transcriptional modifications. **D)** Representative images of *E. coli* biofilms grown in LB medium (top row, +NaCl) and LB medium without salt (bottom row, -NaCl) with all cells constitutively expressing mKOκ, *csgBAC*-expression reflected in mKATE2 levels and CsgA-His labelled with anti-His antibody with Alexa Fluor 488 fluorescence. It was evident that cells expressed *csgBAC* in both conditions, but curli fibres were only present in the bottom row, where cells were grown without salt in the medium. Scale bars represent 10 μm.

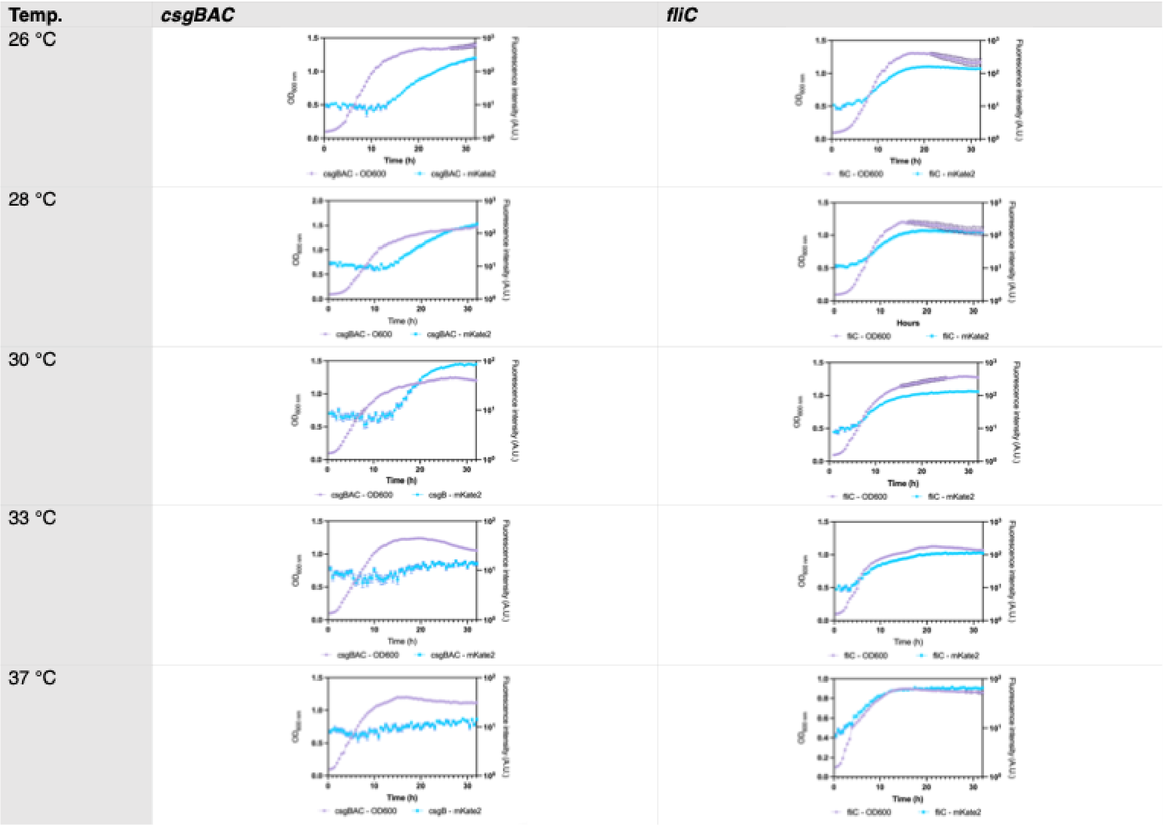


**Supplementary Figure 5) Temporal growth and expression pattern of *csgBAC* at various temperatures.** Expression of *csgBAC* was monitored with a fluorescent transcriptional reporter (*csgBAC*-mKate2) (blue line) in a plate reader also measuring $OD_{600}$ (magenta line). Expression of *csgBAC* increased after 10-15h of growth at 30°C or lower, while not increasing at any point at higher temperatures. A reporter of *fliC* (*fliC*-mKate2) was included as control, as this gene previously have been found to be expressed independent of temperature[2].

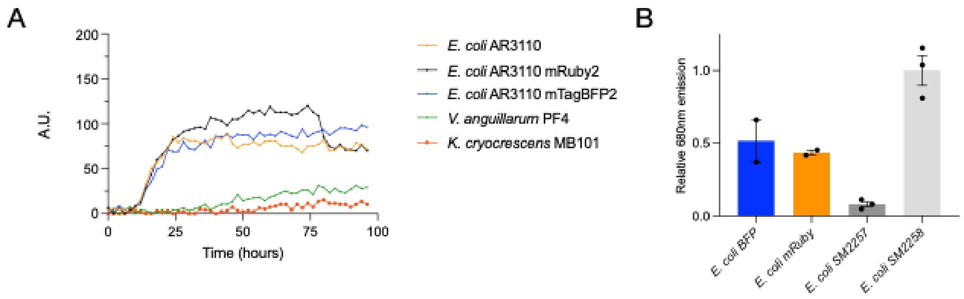


**Supplementary Figure 6) Attesting the ability to form amyloid fibres in colonies. A)** Temporal curli production was quantified with the optotracer EbbaBiolight680, which is a binding-induced fluorescence tracer that previously has been benchmarked as able to quantify amyloid production in *Salmonella enterica* and uropathogenic *E. coli*[1]. The optotracer was used to monitor and quantify curli production over time on LB agar plates w.o. salt at 30°C, where we observed a steep increase in fluorescence after approx. 15 hours for the three *E. coli* strains included (The non-modified wildtype, a mRuby2 variant from a previous study [2] and the mTagBFP2 used in this study), which saturated after approx. 40 hours. The two other species of our model community, *V. anguillarum* and *K. cryocrescens*, were unable to induce fluorescence indicating that they do not form amyloid fibres. **B)** EbbaBiolight680 was also used for endpoint measurement of amyloid fibre production on 72h old colonies, where control strains SM2257 (amyloid neg.) and SM2258 (amyloid pos.) were included as control strains[3]. Again, the mTagBFP2- and mRuby2-expressing strains produce a similar amount of amyloid fibres.

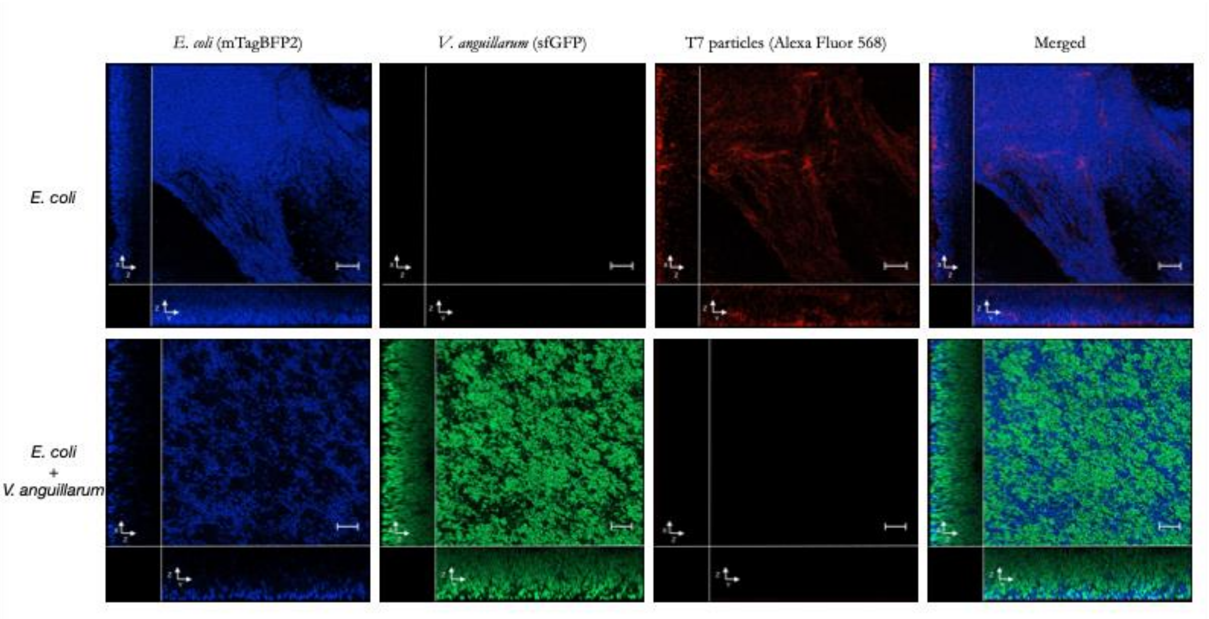


**Supplementary Figure 7) Imaging of phage localization in biofilm communities.** When added to a monospecies biofilm community of *E. coli* (mTagBFP2/blue cells) (top row), labelled T7 (Alexa Fluor 568/red particles) aggregated all over the community, and also reached the inner and bottom part of the biofilm. In contrast, when *E. coli* was co-incubated with *V. anguillarum* on top (sfGFP/green cells) (bottom row), the T7 particles did not attach and were not visible at all on or in the community.

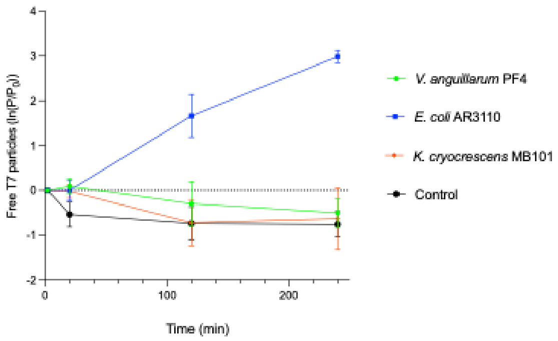


**Supplementary Figure 8) Temporal measurement of free phage particles with different biofilm hosts.** The number of free T7 particles were quantified by plaque assay at various timepoints after being inoculated in wells with 72h old biofilms of *E. coli*, *V. anguillarum* and *K. cryocrescens*, respectively. *E. coli* was the only host able to proliferate the number of T7 particles. The T7 particles were not trapped in the matrix of neither *V. anguillarum* nor *K. cryocrescens*, as the slight decrease in the number of free phages is not exceeding that of the control with a sterile glass slide.

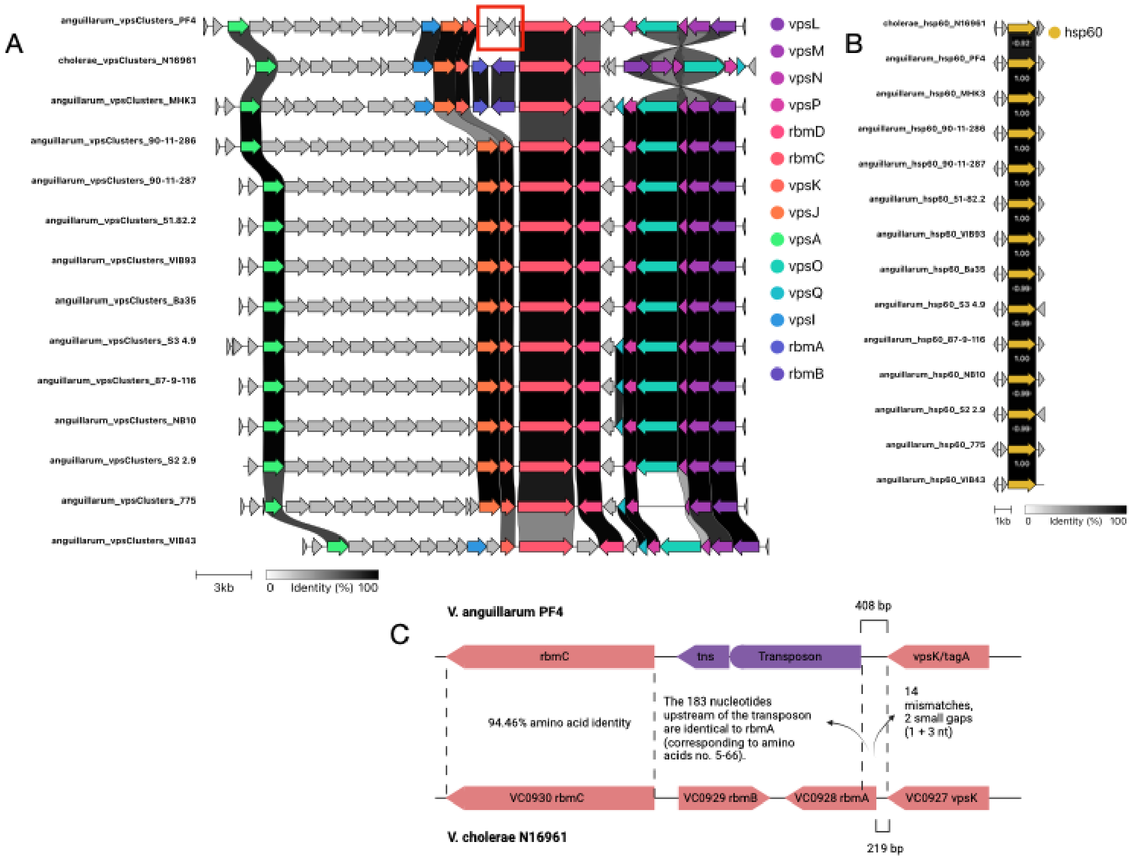


**Supplementary Figure 9) Synteny alignment and comparison of matrix gene clusters. A)** Alignment of the region encoding *vps* and *rbm* genes for available *V. anguillarum* genomes reveal that many of the matrix genes identified in *V. cholerae* (represented by *V. cholerae* N16961, 2nd row) are also present in *V. anguillarum* and with a similar synteny. *rbmA* and *rbmB* is however only present in a single *V. anguillarum* strain (strain MHK3, 3rd row). In the strain used in this study (strain PF4, 1st row) are two hypothetical and a transposase present (marked with red square) at the region where *rbmA* and *rbmB* are located in N16961 and MHK3, which could suggest that these matrix-component encoding genes are generally absent in *V. anguillarum* due to transposon activity. PF4 is however the only strain with these genes located in this region. **B)** To ensure that the *rbmAB*-encoding strain MHK3 were indeed *V. anguillarum*, the gene hsp60 was aligned as well. This housekeeping gene have been found superior to 16S for distinguishing Vibrio species[4,5]. The alignment at amino acid level was minimum 99% identical to the other *V. anguillarum* strains, and only 92% identical to *V. cholerae* N16961 suggesting that MHK3 is likely a *V. anguillarum* strain encoding *rbmAB*. The alignments were made with the Clinker tool software[6]. All accession numbers are available in supplementary table 1. **C)** A closer look at the area with the transposase (tns) (red square) reveal that there are some reminisces of the *rbmA* gene as found in *V. cholerae*; In the upstream region of the transposon, there is nucleotide similarity corresponding to the sequence encoding amino acids number 5 to 66, and some mismatch mutations and deletions upstream of this identical region.

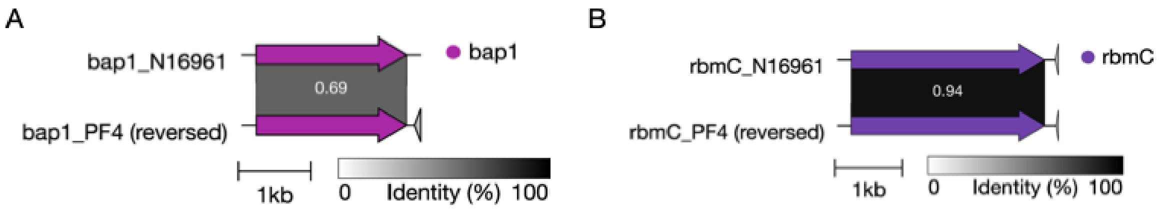


**Supplementary Figure 10) Alignment and comparison of matrix proteins RbmC and Bap1. A)** Alignment of the region encoding rbmC in *V. cholerae* N16961 and *V. anguillarum* PF4. On amino acid level the two proteins are 94% identical. **B)** Alignment of the region encoding bap1 in *V. cholerae* N16961 and *V. anguillarum* PF4. On amino acid level the two proteins are 69% identical. The alignments were made with the Clinker tool software[6].

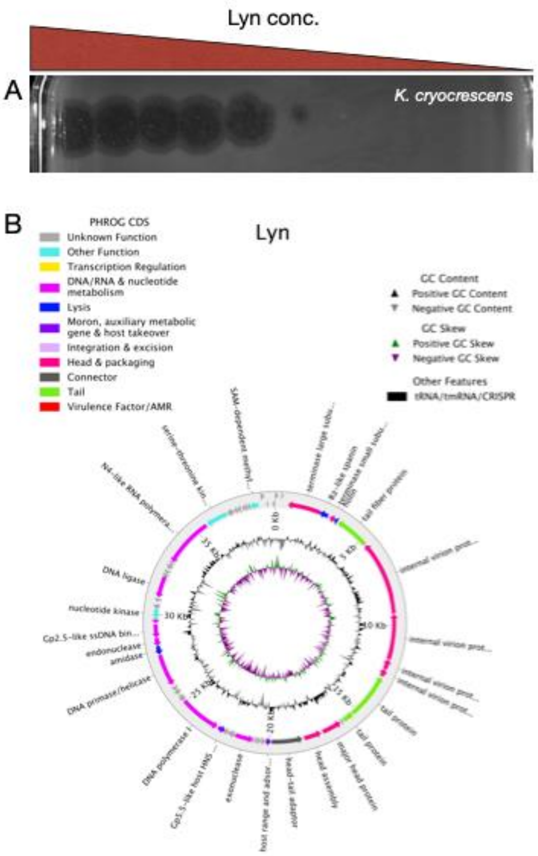


**Supplementary Figure 11) Characterization of kluyveraphage Lyn. A)** Plaque morphology of Lyn at different concentrations. The phage did not form plaques on the two other bacteria, *V. anguillarum* and *E. coli* (not shown). **B)** Genomic characterization and annotation of the 39408-nucleotide genome.

**Supplementary Table 1:** Bacterial strains, bacteriophages, plasmids, reagents and software used in this study

| REAGENT or RESOURCE | SOURCE | IDENTIFIER |
|---|---|---|
| **Antibodies** | | |
| Penta•His Alexa Fluor 488 conjugate | QIAGEN | Cat#35310 |
| **Bacterial and virus strains** | | |
| *E. coli* S17-1 λpir | Simon et al.[7] | Sp1476 |
| *E. coli* AR3110 | Serra et al.[8] | Sp1500 |
| *E. coli* AR3110, CmR, Tn7::Plpp_mTagBFP2 | This study | Sp1535 |
| *E. coli* AR3110, KanR, Tn7::Plpp_mCherry | This study | Sp1632 |
| *E. coli* AR3110, KanR, attB::Ptac_mRuby2 | Vidakovic et al.[2] | KDE679 |
| *E. coli* AR3110, KanR, attB::Ptac_mKOκ, *csgBAC*-mKate2 | Vidakovic et al.[2] | KDE771 |
| *E. coli* AR3110, KanR, attB::Ptac_mKOκ, *csgBAC*-mKate2, *csgA*+6xHIS | Vidakovic et al.[2] | KDE923 |
| *E. coli* AR3110, KanR, attB::Ptac_mKOκ, *fliC*-mKate2 | Vidakovic et al.[2] | KDE782 |
| *E. coli* MG1655 | Lab stock | Sp437 |
| *V. anguillarum* PF4 | Silva-Rubio et al.[9] | Sp1499 |
| *V. anguillarum* PF4, KanR, Tn7::Ptac_sfGFP(Vc) | This study | Sp1531 |
| *V. anguillarum* PF4, KanR, Tn7::Ptac_sfGFP(Vc), Δ*csgD* | This study | Sp1588 |
| *V. anguillarum* PF4, KanR, Tn7::Ptac_sfGFP(Vc), Δ*vpsL* | This study | Sp1592 |
| *V. anguillarum* PF4, KanR, Tn7::Ptac_sfGFP(Vc), Δ*vpsMNOP* | This study | Sp1596 |
| *V. anguillarum* PF4, KanR, Tn7::Ptac_sfGFP(Vc), Δ*rbmC* | This study | Sp1668 |
| *V. anguillarum* PF4, KanR, Tn7::Ptac_sfGFP(Vc), Δ*bap1* | This study | Sp1682 |
| *V. anguillarum* PF4, KanR, Tn7::Ptac_sfGFP(Vc), Δ*vpsIJ* | This study | Sp1693 |
| *K. cryocrescens* MB101 | Burmølle et al.[10] | Sp647 |
| *K. cryocrescens* MB101, KanR, Tn7::Plpp_mCherry | This study | Sp1533 |
| Coliphage T7 | Demerec & Fano[11] | Sp1539 |
| Kluyveraphage Lyn | This study | Sp1785 |
| **Chemicals, peptides, and recombinant proteins** | | |
| Sodium chloride | VWR | Cat#27810.295 |
| Yeast extract | VWR | Cat#J850-500G |
| Granulated tryptone | Millipore | Cat#1.07213.1000 |
| Difco agar noble | BD | Cat#214230 |
| Agar-agar | VWR | Cat#J637-2.5kg |
| TCBS-agar | Millipore | Cat#86348-500G |
| EbbaBiologht 680 | Ebba Biotech | Cat#Ebba680 |
| PEG8000 | Promega | Cat#V3011 |
| Tris-HCl | Roche | Cat#10812846001 |
| Gelatin | Sigma-Aldrich | Cat#G1890-100G |
| $MgSO_4 \bullet 7H_2O$ | VWR | Cat#25167.298 |
| DNase | A&A Biotechnology | Cat#1009-100 |
| Proteinase K | A&A Biotechnology | Cat#1019-20 |
| **Critical commercial assays** | | |
| Alexa Fluor 568 labelling kit | ThermoFisher | Cat#A10238 |
| Phusion Hot Start II DNA polymerases | ThermoFisher | Cat#F549S |
| PCRBIO Taq DNA polymerases | PCRBiosystems | Cat#PB10.11-05 |
| QIAEX II Gel Extraction kt | QIAGEN | Cat#20021 |
| QIAquick PCR purification kit | QIAGEN | Cat#28104 |
| DNA Clean & Concentrator-5 | Zymo Research | Cat#D4003 |
| DNeasy UltraClean Microbial Kit | QIAGEN | Cat#12224.50 |

| | | |
|---|---|---|
| NEBNext Ultra II FS Library prep kit | New England Biolabs | Cat#E7805L |
| Nextera XT DNA library prep kit | Illumina | Cat#FC-131-1096 |
| Rapid barcoding kit | Nanopore | Cat#SQK-RBK110.96 |
| NEBuilder HiFi DNA Assembly Master mix | New England Biolabs | Cat#E2621X |
| **Oligonucleotides** | | |
| Supplementary table 2 | TAG Copenhagen | N/A |
| **Recombinant DNA** | | |
| Plasmid pDM4 (in *E. coli* S17-1 λpir) | Milton et al.[12] | Sp1475 |
| Plasmid pNUT542 (in *E. coli* TOP10) | Singh et al.[13] | KDE542 |
| Plasmid pUC18R6KT-mini-Tn7T-Km (in *E. coli* pir1) | Choi et al.[14] | KDE2658 |
| Plasmid pNUT2703 (in *E. coli* S17-1 λpir) | This study | KDE2703 |
| Plasmid pRK2013 (in *E. coli* S17-1 λpir) | Ditta el al.[15] | Sp1470 |
| Plasmid pTN2 (in *E. coli* pir1) | Choi et al.[14] | Sp1469 |
| Plasmid pGRG36_Plpp_mCherry_KanR (in *E. coli* S17-1 λpir) | Olesen et al.[16] | Sp1310 |
| Plasmid pNUT2787 (in *E. coli* S17-1 λpir) | Diaz-Pascual et al.[17] | KDE2787 |
| Plasmid pmTagBFP2-pBAD (in *E. coli* DH5α) | Subach et al.[18] | Sp1397 |
| Plasmid pMFH7 (in *E. coli* S17-1 λpir) | This study | Sp1529 |
| Plasmid pMFH63 (in *E. coli* S17-1 λpir) | This study | Sp1582 |
| Plasmid pMFH78 (in *E. coli* S17-1 λpir) | This study | Sp1624 |
| Plasmid pMFH79 (in *E. coli* S17-1 λpir) | This study | Sp1625 |
| Plasmid pJHO64 (in *E. coli* S17-1 λpir) | This study | Sp1584 |
| Plasmid pJHO65 (in *E. coli* S17-1 λpir) | This study | Sp1583 |
| Plasmid pJHO66 (in *E. coli* S17-1 λpir) | This study | Sp1581 |
| **Software and algorithms** | | |
| MATLAB | Mathworks | v.R2023b |
| BiofilmQ | Hartmann et al.[19] | v.0.2.2 |
| Snapgene | Dotmatics | v.8.1 |
| Graphpad Prism | Software MacKiev | v.10.3.0 |
| CLC Genomics Workbench | QIAGEN | v.12.0.3 |
| TYGS | Meier-Kolthoff et al.[20] | N/A |
| Clinker | Gilchrist & Chooi[6] | N/A |
| SPAdes | Bankevich et al.[21] | v.3.13.1 |
| taxMyPhage | Millard et al.[22] | N/A |
| pharokka | Bouras et al.[23] | N/A |
| FastP | Chen et al.[24] | v.0.23.4 |
| Unicycler | Wick et al.[25] | v.0.4.8 |
| Guppy | Wick et al.[26] | v.5.1.13 |
| Filtlong | Github.com/rrwick/Filtlong | v.0.2.1 |
| Flye | Kolmogorov et al.[27] | v.2.9.1 |
| CheckM | Parks et al.[28] | N/A |
| **Other** | | |
| 12-well culture plates | Greiner Bio-One | Cat#665 180 |
| Ø15mm round cover glasses | Marienfeld | Cat#0111550 |
| 24x50mm cover glass #1.5 | Thorlabs | Cat#CG15KH |
| Luer lock syringe 50ml | VWR | Cat#613-2053 |
| Syringe filters, 0.22µm PVDF | Carl Roth | Cat#P666.1 |
| Syringe filters, 0.45µm PVDF | Carl Roth | Cat#P667.1 |
| MCE membrane filters, 0.22µm | Millipore | Cat#GSWP09000 |

**Supplementary Table 2:** Genome accession numbers for comparison of matrix-encoding gene organization in *V. anguillarum.*

| Strain | Genome accession no. |
|---|---|
| V. anguillarum NB10 | LK021130 |
| V. anguillarum MKH3 | CP022468 |
| V. anguillarum 1-1(7) | Not assembled – contigs GCA_030851405.1 |
| V. anguillarum 850617-1/1 | Not assembled – scaffold GCA_015350525.1 |
| V. anguillarum 040915-1/1B | Not assembled – Scaffold GCA_015343015.1 |
| V. anguillarum 775 | CP002284 |
| V. anguillarum VIB43 | CP023054 |
| V. anguillarum 87-9-116 | CP021980 |
| V. anguillarum PF4 | CP010080 |
| V. anguillarum 90-11-287 | CP011475 |
| V. anguillarum 51/82/2 | CP010042 (partial) |
| V. anguillarum S3 4/9 | CP022099 |
| V. anguillarum S2 2/9 | CP011472 (partial) |
| V. anguillarum VIB93 | CP011438 (partial) |
| V. anguillarum Ba35 | CP010030 (partial) |
| V. anguillarum 90-11-286 | CP011460 |
| V. cholerae N16961 | LT906614 |

**Supplementary Table 3:** Oligos used in this study

| Name | 5'-Sequence-3' | Description |
|---|---|---|
| MFHO170 | TCGACGGTATCGATAAGCTTG | Fw: Amplification of pDM4 as backbone |
| MFHO171 | TGGGGCCCTTCTAGATAGAT | Rv: Amplification of pDM4 as backbone |
| MFHO166 | ATCTAGAAGGGCCCCATCGATGCGAACACACACAC | Fw: amplification of upstream fragment flanking vpsL |
| MFHO167 | CTAGTAAGCTTCACCCATTGTAGAAGCTCCT | Rv: amplification of upstream fragment flanking vpsL |
| MFHO168 | CAATGGGTGAAGCTTACTAGAATGGATAAAGCATACTC | Fw: amplification of downstream fragment flanking vpsL |
| MFHO169 | GCTTATCGATACCGTCGAGTGCTGGTGTAATTCTCGC | Rv: amplification of downstream fragment flanking vpsL |
| MFHO172 | ATCTAGAAGGGCCCCACGATGAGCGTAACAAAAGCT | Fw: amplification of upstream fragment flanking csgD |
| MFHO173 | CACATAATTAAGACCTCATGGTCACTTCCTGTTG | Rv: amplification of upstream fragment flanking csgD |
| MFHO174 | ACCATGAGGTCTTAATTATGTGCTTTCACGC | Fw: amplification of downstream fragment flanking csgD |
| MFHO175 | GCTTATCGATACCGTCGACGCTAGGTGACTAACGCA | Rv: amplification of downstream fragment flanking csgD |
| MFHO176 | CGAAGTGATCCATGATCG | Fw: verification of csgD deletion |
| MFHO177 | CCATGAATCTGACGTTGC | Rv: verification of csgD deletion |
| MFHO178 | CACGATATAGCGTGACTTACC | Fw: verification of vpsL deletion |
| MFHO179 | GTTGTCGTTATAACCCGTTG | Rv: verification of vpsL deletion |
| MFHO180 | ATCTAGAAGGGCCCCAGCTGAAAGCCTTGATATATACA | Fw: amplification of upstream fragment flanking vpsMNOP |
| MFHO181 | CTTTCATATGAGTATGCTTTATCC | Rv: amplification of upstream fragment flanking vpsMNOP |
| MFHO182 | GGATAAAGCATACTCATATGAAAGAGTGATCTCGATGATA AAACA | Fw: amplification of downstream fragment flanking vpsMNOP |
| MFHO183 | GCTTATCGATACCGTCGAGGGCAGGAATATACCTGC | Rv: amplification of downstream fragment flanking vpsMNOP |

| | | |
|---|---|---|
| MFHO184 | ATCTAGAAGGGCCCCATGCAGCCCTTATATATCAACGA | Fw: amplification of upstream fragment flanking vpsJI |
| MFHO185 | GAATAACCGATTGAAACGC | Rv: amplification of upstream fragment flanking vpsJI |
| MFHO186 | TGCGTTTCAATCGGTTATTCGGCGACCATTAGATTAATCG | Fw: amplification of downstream fragment flanking vpsJI |
| MFHO187 | GCTTATCGATACCGTCGAAGGCTAATGAGTTTTCGC | Rv: amplification of downstream fragment flanking vpsJI |
| MFHO188 | GTGTGAGCTATCGAGTTTTGC | Fw: verification of vpsMNOP deletion |
| MFHO189 | GCAATGAATCATAAACCCACACG | Rv: verification of vpsMNOP deletion |
| MFHO190 | CGCTTTCACGTAAAATGGTCAG | Fw: verification of vpsJI deletion |
| MFHO191 | CACACGAGACAAAGTATCTGC | Rv: verification of vpsJI deletion |
| MFHO192 | CGAACTAAACCCTCATGG | Fw: Verification of insertion in pDM4-derivates |
| MFHO193 | CTCAAAAAATACGCCCGG | Rv: Verification of insertion in pDM4-derivates |
| MFHO244 | CGACTTGAAGTTGCAGG | Fw: verification of rbmC deletion |
| MFHO245 | GATGTCTCATTAGTGAGGC | Rv: verification of rbmC deletion |
| MFHO246 | CTATCTAGAAGGGCCCCAGAAATCTGGATTGCATTG | Fw: amplification of downstream fragment flanking rbmC |
| MFHO247 | CTTACAGATGACGGTATTCGATTAACTGATATCG | Rv: amplification of downstream fragment flanking rbmC |
| MFHO248 | CGAATACCGTCATCTGTAAGACTTCC | Fw: amplification of upstream fragment flanking rbmC |
| MFHO249 | GCTTATCGATACCGTCGACGTGGTTCTTATGATCGC | Rv: amplification of upstream fragment flanking rbmC |
| MFHO250 | GTTGGCCGTATTAATCGTG | Fw: verification of bap1 deletion |
| MFHO251 | GACCTATCGTTGAAGTGCA | Rv: verification of bap1 deletion |
| MFHO252 | GCTTATCGATACCGTCGAGGTTAGACTTCTGCAATCAC | Fw: amplification of downstream fragment flanking bap1 |
| MFHO253 | GTACTTGAGCGAGAAGTCATTAGGTAACATCTCTC | Rv: amplification of downstream fragment flanking bap1 |

| | | |
|---|---|---|
| MFHO254 | CTATCTAGAAGGGCCCCAGAGCTCGCGACTAAACTTG | Fw: amplification of upstream fragment flanking bap1 |
| MFHO255 | GACTTCTCGCTCAAGTACCATTAAATAACTTG | Rv: amplification of upstream fragment flanking bap1 |
| KDO3817 | TCGACTCCTTCTCGAGGAATTCCTGCAGCC | Fw: amplification of pUC18R6KT-mini-Tn7T-Km |
| KDO3818 | GAATGCTGGTACCTCGCGAAGGCCT | Rv: amplification of pUC18R6KT-mini-Tn7T-Km |
| KDO3913 | CGCGAGGTACCAGCATTCCATTTCACACCTCCTGTACGC | Fw: amplification of Ptac_sfGFP(Vc) from pNUT542 |
| KDO3914 | TTCCTCGAGAAGGAGTCGAACCGGATCCGCTAGCAGG | Rv: amplification of Ptac_sfGFP(Vc) from pNUT542 |
| MFHO17 | CGCGAGGTACCAGCATTCCGATGAATCCGATGGAAGCA | Fw: amplification of Plpp promoter from pGRG36_Plpp_mCherry_KanR |
| MFHO18 | CATCAACCACCTCCTATACCCTCTAGATTGAGTTAATCTCC | Rv: amplification of Plpp promoter from pGRG36_Plpp_mCherry_KanR |
| MFHO19 | GTATAGGAGGTGGTTGATGGTGTCTAAGGGCGAAG | Fw: amplification of mTagBFP2 from mTagBFP2-pBAD |
| MFHO20 | TTCCTCGAGAAGGAGTCGACTTCGAATTCTTAATTAAGCTTGTGC | Rv: amplification of mTagBFP2 from mTagBFP2-pBAD |
| MFHO9 | CACCGATCTTCTACACCGTTCCGC | Fw: verification of Tn7 insertion in *E. coli* |
| MFHO10 | AGATCAGTTTGGTGTACGCCAGGT | Rv: verification of Tn7 insertion in *E. coli* |
| MFHO69 | GATGCTGGTGGCGAAGCTGT | Fw: verification of Tn7 insertion in *K. cryocrescens* |
| MFHO70 | GATGACGGTTTGTCACATGGA | Rv: verification of Tn7 insertion in *K. cryocrescens* |
